# PhyloZoo: a unified framework for phylogenetic network analysis in Python

**DOI:** 10.64898/2026.06.09.731120

**Authors:** Niels Holtgrefe

**Affiliations:** Delft Institute of Applied Mathematics, Delft University of Technology, Mekelweg 4, 2628 CD, Delft, The Netherlands

## Abstract

Reticulate evolutionary processes (events in which lineages merge, such as hybridization, recombination, and horizontal gene transfer) are widespread across nature but cannot be represented by phylogenetic trees alone. Phylogenetic networks have therefore become an important modelling tool, yet existing software is typically tied to specific inference paradigms and provides limited support for working with multiple network representations in a unified and programmable environment.

PhyloZoo is an open-source Python framework that lowers the barrier to developing practical, easy-to-use software for phylogenetic network analysis. It provides data structures and algorithms covering the main representations used in the field, together with dedicated visualization tools and robust I/O for all major phylogenetic file formats. A particular emphasis lies on semi-directed phylogenetic networks, which explicitly represent root uncertainty and have so far received limited support in existing software. By offering a shared, scalable foundation for developing interoperable tools and a combinatorial layer that supports computational proofs and theoretical exploration, PhyloZoo enables reproducible workflows for applied, methodological, and theoretical studies of reticulate evolution.

**Availability and implementation:** PhyloZoo is implemented in Python and installable from PyPI, with source code, documentation, and examples available at https://github.com/nholtgrefe/phylozoo.

## 1. Introduction

Reticulate evolutionary events, such as hybridization, horizontal gene transfer, and recombination, are recognized as pervasive across microbial and eukaryotic lineages (Huson et al., 2010; Kong et al., 2022; Soucy et al., 2015). Because phylogenetic trees cannot represent the merging of lineages, phylogenetic networks have become a central modelling tool in evolutionary biology, extending the tree model to accommodate taxa whose ancestry derives from multiple parental lineages.

A realistic network analysis workflow draws on a diverse set of representations (Huson et al., 2010). Starting from *sequence alignments*, a researcher may compute pairwise *distance matrices*, summarize gene-tree collections as *triplet* (rooted three-taxon topology) or *quartet* (unrooted four-taxon topology) data, or extract *split systems* that represent bi-partitions of taxa. Then, they may represent the data as an *implicit*, undirected split network, or infer an *explicit* phylogenetic network, which can either be *directed* or *semi-directed*. The latter is a mixed graph that is increasingly popular because it captures root uncertainty (Solís-Lemus and Ané, 2016). Moving reliably between these representations requires sustained manual effort when no unified software environment exists.

PhyloZoo is an open-source Python package that brings together these representations within a single environment, designed to serve a broad spectrum of users in phylogenetic network analysis. It offers robust I/O functionality and rich conversions between validated classes for several major datatypes, and its core operations scale to networks with 10^5^ leaves and 10^3^ reticulations (see Table 1). The package provides dedicated visualization tools for directed and semi-directed networks (see Figure 1), which further enable the development of graphical interfaces and accessible user applications. The packages Squirrel (Holtgrefe et al., 2025) and PhyloCass (van Iersel, 2026) (both for network reconstruction) and PaNDA (Holtgrefe et al., 2026) (for phylogenetic diversity computation), demonstrate how shared PhyloZoo infrastructure enables efficient development of interoperable tools. This reusable foundation frees *method developers* to focus on algorithmic novelty without managing low-level infrastructure and I/O functionality. Furthermore, *theorists* gain programmatic access to combinatorial concepts—including network class checking, isomorphism testing, and structures such as split systems, triplets, quartets, and generators— which would otherwise be tedious to manipulate by hand, facilitating rigorous theoretical prototyping and exploration.

**Table 1.** Runtimes (in seconds) of fundamental PhyloZoo operations. For each size, 20 directed networks were generated by uniformly adding *r* reticulation edges to a balanced tree with *n* leaves; each operation was timed three times per network on the directed network (D) or on its converted semi-directed network (SD), and the average is reported. Subnetworks were extracted on *n/*2 leaves.

| Operation | $n$ leaves, $r$ reticulations | | |
| --- | --- | --- | --- |
| | $10^3, 10$ | $10^4, 10^2$ | $10^5, 10^3$ |
| Loading from eNewick (D) | 0.02 | 0.27 | 3.3 |
| Extracting a subnetwork (D) | 0.01 | 0.16 | 1.9 |
| Converting D to SD | 0.02 | 0.22 | 3.0 |
| Validating the network (SD) | 0.04 | 0.51 | 6.0 |
| Saving as eNewick (D) | 0.004 | 0.04 | 0.6 |

All timings are single-threaded, measured on a laptop with an Intel Core i5-1245U (up to 4.4 GHz) and 16 GB RAM, running Debian 12 (Linux 6.1) and Python 3.11.

**Figure 1.**
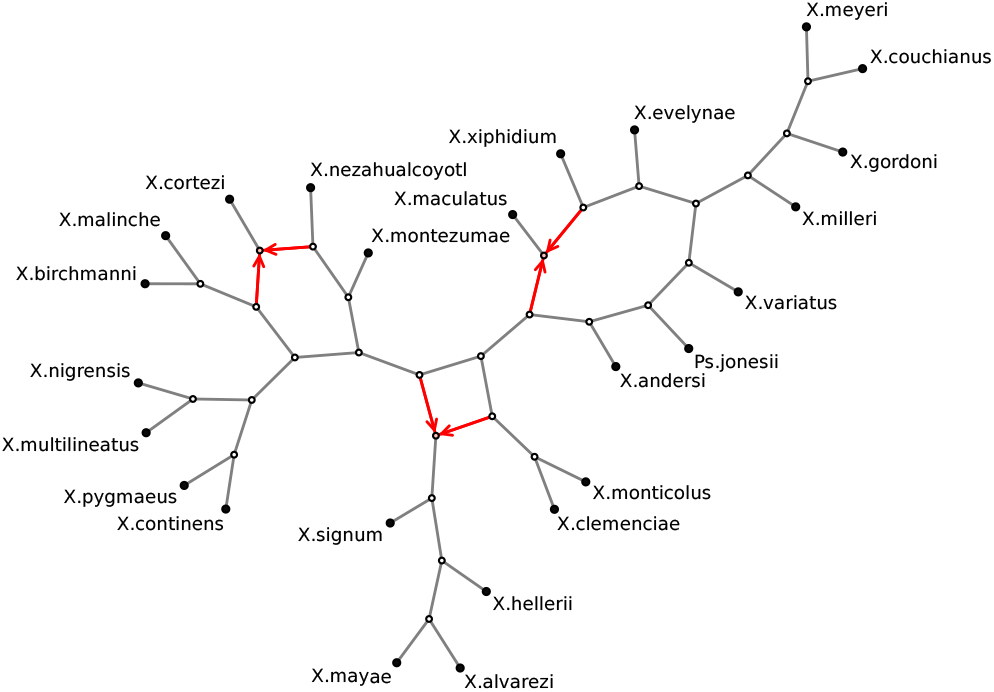
A semi-directed phylogenetic network on *Xiphophorus* fishes, inferred with Squirrel (Holtgrefe et al., 2025) and visualized using PhyloZoo. Reticulation edges are highlighted in red.

### Related software

Mature frameworks exist for phylogenetic *trees*, notably phangorn (Schliep, 2011) and ape (Paradis and Schliep, 2019) in R, and DendroPy (Sukumaran and Holder, 2010) and ETE3 (Huerta-Cepas et al., 2016) in Python. Some extend beyond trees, though only secondarily: phangorn handles implicit split networks but not explicit ones, while ape does allow directed networks, but only basic storing and plotting (with the recent R package tanggle (Schliep et al., 2026) providing improved plotting for both packages). Software dedicated primarily to networks is more fragmented: much of it consists of single-purpose scripts that are not easily combined. Among the more substantial frameworks, graphical interfaces such as Split-sTree App (Huson and Bryant, 2024) excel at displaying and exploring data but offer limited programmability. PhyloX (Janssen, 2024) (Python) offers a specialized set of utilities for directed networks, whereas PhyloNetworks (Solís-Lemus et al., 2017) (Julia) provides a broader programmable foundation including network data structures, visualization, and utility functions. Other packages, such as MSCquartets (Rhodes et al., 2021) (R) and the Java tools BEAST (Baele et al., 2025) and PhyloNet (Than et al., 2008), provide reusable functionality but are structured around specific statistical inference paradigms. Unlike these, PhyloZoo adopts a combinatorial stance, bringing directed and semi-directed networks, split systems, triplet and quartet data, and network generators together in a single, programmable Python environment.

## 2. Core features

PhyloZoo is organized into three modules: `core` (data structures and algorithms), `viz` (visualization), and `utils` (I/O and validation). All major classes are validated after construction, ensuring objects always represent well-formed phylogenetic structures; validation may be disabled for performance-critical use cases. Figure 2 gives an impression of core datatypes, derived structures, and functions available in the package.

**Figure 2.**
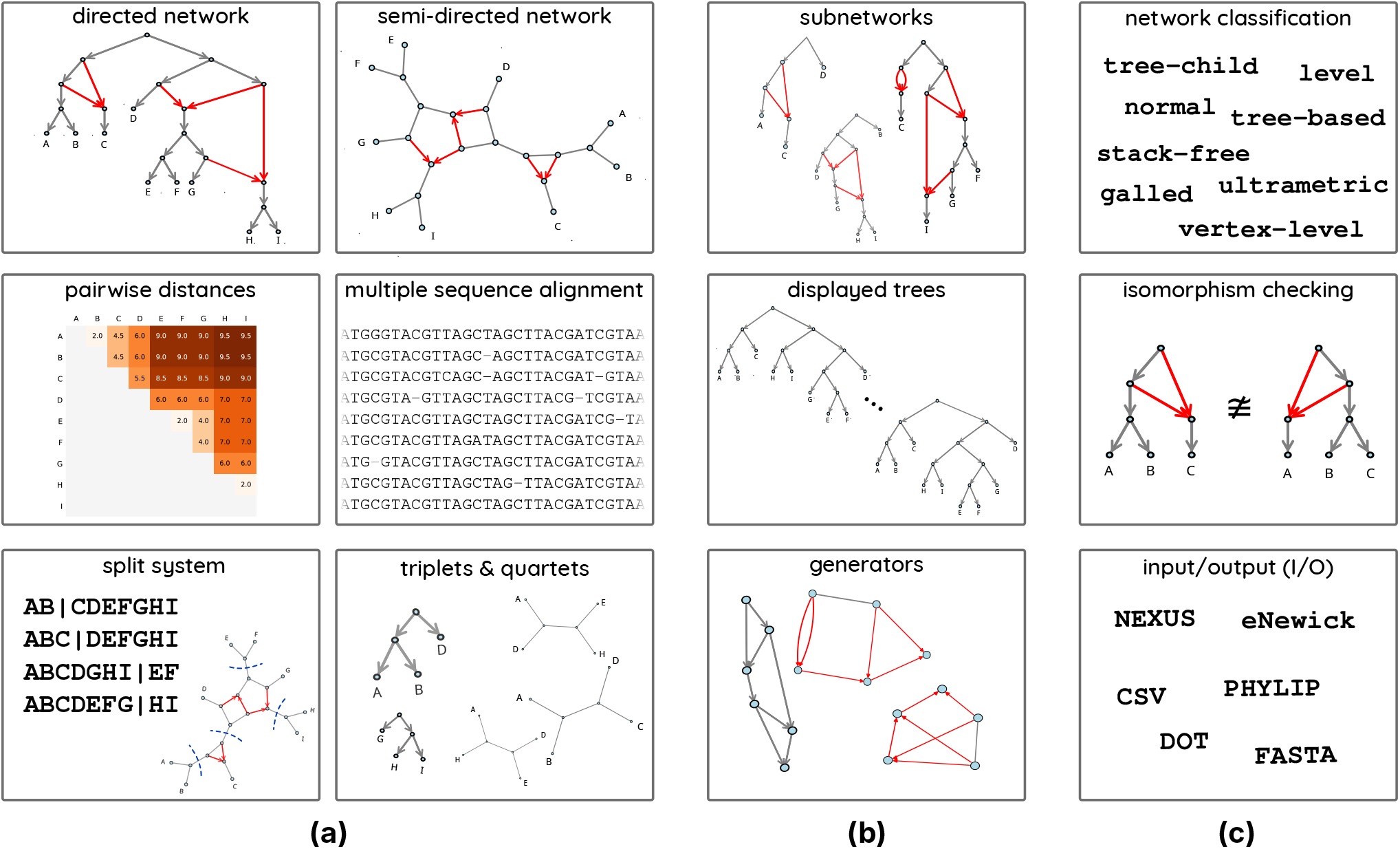
Overview of PhyloZoo’s core functionality: supported datatypes (a), structures derived from network objects (b), and key operations (c).

### Phylogenetic networks

The two central classes represent *explicit* phylogenetic networks, in either *directed* or *semi-directed* form. A directed (phylogenetic) network is a directed acyclic graph with an explicit root, internal tree nodes, reticulation nodes, and labelled leaves. A semi-directed (phylogenetic) network does not have a root and leaves all non-reticulation edges undirected, so that only edges incident to reticulation nodes retain an orientation; this models root uncertainty while preserving reticulation information (see Figure 1). Both classes support non-binary nodes, parallel edges, branch lengths, bootstrap support values, and inheritance probabilities, with dedicated routines enabling conversion between them.

A rich set of structural operations is available for both network types (see Figure 2(b,c)). Membership in structural network classes (which constrain the placement of reticulations, including tree-child, tree-based, galled, stack-free, and normal; see also (Kong et al., 2022)) can be tested for both directed and semi-directed representations. Derived objects, such as displayed trees, split systems, triplets, quartets, subnetworks, and pairwise distances are all directly extractable from a network, and isomorphism checking is supported via NetworkX (Hagberg et al., 2008).

PhyloZoo also includes support for network *generators* (Gambette et al., 2009)—the minimal structural skeletons from which all networks with a given number of reticulations are constructed by attaching leaves—for both directed and semi-directed representations. Generators can be exhaustively enumerated up to isomorphism and used to construct or benchmark complete sets of network topologies, supporting both the evaluation of algorithms and systematic theoretical exploration (see Section 3).

#### Related datatypes

Beyond networks, PhyloZoo provides validated classes for several major datatypes used in phylogenetics. See Figure 2(a) for illustrations.

*Multiple sequence alignments* store aligned sequences with array-backed representations. Both bootstrapping and direct computation of pairwise distances are supported.

*Pairwise distances* between taxa can be stored in a dedicated distance matrix class, which supports classification checks (metric, circular and totally decomposable). Distances can also be computed directly from a network object as shortest, longest, or average path lengths between taxa.

*Split systems* represent weighted or unweighted bipartitions of the taxon set, a standard summary of the phylogenetic signal in a data set. They include algorithms for tree reconstruction, distance computation, quartet extraction, and split decomposition.

*Triplet and quartet data* (rooted three-taxon and unrooted four-taxon subtree topologies, respectively) can be extracted from networks and compared across datasets, with optional weights to represent confidence, such as for quartet concordance factors (Solís-Lemus and Ané, 2016).

### Visualization

A distinguishing contribution of PhyloZoo is its visualization suite for directed and semidirected networks (see Figures 1 and 2). To our knowledge, it is the first Python package to provide plotting of phylogenetic networks, including layout algorithms and styling options for both network types. The resulting plots integrate directly with standard scientific Python workflows and can be embedded in graphical user interfaces, making PhyloZoo a natural backend for interactive phylogenetic applications.

### Input/Output

PhyloZoo supports reading and writing all major phylogenetic file formats via a uniform load/save interface on every datatype class. Supported formats include extended Newick (eNewick), NEXUS, FASTA, PHYLIP, DOT, and CSV. A *format registry* allows developers to register custom file format parsers with minimal additional code, making the package straightforward to extend as new formats emerge.

### Performance

Table 1 reports runtimes for fundamental operations on networks of increasing size and complexity, illustrating the order of magnitude that can be expected in practice. Even for networks with 10^5^ leaves and 10^3^ reticulations, loading and saving extended Newick, validating, converting a directed network into a semi-directed one, and extracting a subnetwork all complete within seconds. Such operations are therefore not the bottleneck when building algorithms on top of PhyloZoo: what dominates is the runtime of the algorithm itself. Operations that are algorithmically heavy naturally scale differently; enumerating the displayed trees of a network, for example, takes time exponential in the number of reticulations *r* and is thus already infeasible for *r* = 10^2^.

## 3. Applications

### Network reconstruction

Squirrel (Holtgrefe et al., 2025) is a package that infers level-1 semi-directed networks from multi-locus sequence data. Because Squirrel builds on PhyloZoo, it can exploit its data structures, I/O and visualization tools: inferred networks are directly plottable, saveable in any supported format, and ready for downstream analyses without reformatting. An example output is shown in Figure 1.

Similarly, PhyloZoo enabled the development of PhyloCass (van Iersel, 2026), a standalone reimplementation of the Cass algorithm (van Iersel et al., 2010) for building a directed network from a set of conflicting trees.

### Phylogenetic diversity computations

PaNDA (Holtgrefe et al., 2026) is a package that generalizes classical phylogenetic diversity measures to phylogenetic networks and implements diversity-maximizing taxon selection algorithms. Because PaNDA operates on the same PhyloZoo objects as Squirrel, the two packages interoperate without data conversion, enabling end-to-end pipelines from sequence alignment to diversity analysis.

### Combinatorial exploration

A central question in phylogenetic network theory is *distinguishability* : whether data such as gene trees, triplets, quartets, or split systems uniquely determine the underlying network, or whether two distinct evolutionary histories could produce identical observations and therefore be impossible to distinguish from such data alone (Kong et al., 2022). Probing such questions may involve exhaustive case analyses over complete sets of network topologies, which are impractical by hand. PhyloZoo’s combinatorial infrastructure supports such theoretical investigations at scale: by combining generator enumeration, displayed-tree extraction, and isomorphism checking, one can systematically search for counterexamples to such encoding questions across complete sets of network topologies. For example, it is known that two distinct networks can display identical sets of gene trees—a fundamental obstacle to identifiability from genomic data under models without incomplete lineage sorting (Pardi and Scornavacca, 2015)—and this can be discovered automatically with PhyloZoo (see the documentation tutorial).

## 4. Conclusion

PhyloZoo provides a unified Python framework for phylogenetic network analysis, integrating validated, scalable data structures, rich interconversions, dedicated visualization, and a clean extensible API. We have demonstrated how it supports the development of easy-to-use tools for network inference and biodiversity conservation, as well as the systematic case-checking that underpins fundamental mathematical work in phylogenetic network research. Together, these examples illustrate the key design benefit of PhyloZoo: a shared, validated foundation on which independent tools and analyses compose naturally. By reducing the infrastructure burden for method developers and theorists alike, and by enabling the construction of accessible graphical tools for applied users, it fills a gap that inference- and visualization-focused packages leave open.

## Acknowledgements

The author thanks Leo van Iersel and Vincent Moulton for valuable discussions leading to the development of this package, and Tim Holtgrefe, Leo van Iersel, and the anonymous reviewers for helpful comments on the manuscript.

## Data availability

The PhyloZoo source code, accompanied by comprehensive documentation and tutorials, is freely available at https://github.com/nholtgrefe/phylozoo, and the package is installable via pip from PyPI. Scripts and data reproducing the benchmarks in Table 1, the network visualization in Figure 1, and the combinatorial exploration in Section 3 are archived on Zenodo (https://doi.org/10.5281/zenodo.22743136), together with version 0.3.0 of the source code, which was used to produce these results.

